# Variation among intact tissue samples reveals the core transcriptional features of human CNS cell classes

**DOI:** 10.1101/265397

**Authors:** Kevin W. Kelley, Hiromi Inoue, Anna V. Molofsky, Michael C. Oldham

**Affiliations:** Dept. of Neurological Surgery, UCSF; Eli and Edythe Broad Center of Regeneration Medicine and Stem Cell Research, UCSF; Dept. of Psychiatry, UCSF; Medical Scientist Training Program and Neuroscience Graduate Program, UCSF

## Abstract

It is widely assumed that cells must be physically isolated to study their molecular profiles. However, intact tissue samples naturally exhibit variation in cellular composition, which drives covariation of cell-class-specific molecular features. By analyzing transcriptional covariation in 7221 intact CNS samples from 840 individuals representing billions of cells, we reveal the core transcriptional identities of major CNS cell classes in humans. By modeling intact CNS transcriptomes as a function of variation in cellular composition, we identify cell-class-specific transcriptional differences in Alzheimer’s disease, among brain regions, and between species. Among these, we show that *PMP2* is expressed by human but not mouse astrocytes and significantly increases mouse astrocyte size upon ectopic expression in vivo, causing them to more closely resemble their human counterparts. Our work is available as an online resource (http://oldhamlab.ctec.ucsf.edu) and provides a generalizable strategy for determining the core molecular features of cellular identity in intact biological systems.

## INTRODUCTION

Identifying the molecular features that define cellular identities is a fundamental goal of biological research. Consequently, a number of methods have been developed to isolate cells for molecular profiling, including fluorescence-activated cell sorting (FACS), immunopanning (IP), and sorting of single cells (SC) or nuclei (SN). Although these methods are readily applied to many biological systems, their applicability to the adult human CNS has been limited by technical factors and practical considerations. For example, FACS, IP, and SC typically require fresh tissue and have therefore been mostly limited to surgical samples from a handful of CNS regions and individuals^1-3^. SN^4, 5^ is compatible with frozen tissue but, like SC, suffers from technical noise caused by tissue dissociation, nucleus/cell capture, cDNA preamplification, and stochastic transcript coverage^6^. Furthermore, there is a trade-off between sequencing depth and the number of nuclei/cells that can be analyzed.

The adult human CNS is large, heterogeneous, and difficult to dissociate due to extensive myelin. It consists of ~170 billion cells, about half of which are neurons^7^. The remaining cells consist mostly of oligodendrocytes, astrocytes, and microglia, which are collectively referred to as glia. Identifying transcriptional differences among neuronal and glial subtypes is an important goal, since the extent of heterogeneity among major CNS cell classes is not fully understood. However, overlooked in the focus on heterogeneity is the equally important question of what CNS cell subtypes have in common. For example, is there a core set of genes whose expression is shared among all neurons? All astrocytes? Etc. Answering these questions will fill critical gaps in our understanding of CNS cell biology, produce novel experimental and analytical strategies, and provide important insights into the cellular origins of CNS pathologies. ‘Bottom-up’ methods such as SC/SN are poorly suited to address these questions, since they are difficult to apply to the adult human CNS at scale.

Most gene expression studies of the human CNS have analyzed intact postmortem tissue samples. Because these samples are heterogeneous and cells must be destroyed to extract RNA, it is often assumed that these datasets contain no information about the cellular origins of gene expression. However, it is axiomatic that intact tissue samples from any biological system will exhibit variation in cellular composition. Therefore, when many intact tissue samples are analyzed, genes expressed with the greatest sensitivity and specificity in the same cell class should appear highly correlated, since their expression levels depend primarily on the proportion of that cell class in each sample^8^. In support of this reasoning, we previously discovered highly reproducible gene coexpression modules in microarray data from intact human brain samples that were significantly enriched with markers of major CNS cell classes^9^. These findings were replicated in studies of intact CNS transcriptomes from mice^10^, rats^11^, zebra finches^12^, macaques^13^, and humans^14^.

Gene coexpression modules corresponding to major cell classes are therefore robust and predictable features of CNS transcriptomes derived from intact tissue samples. Furthermore, the same genes consistently show the strongest affinities for these modules, offering substantial information about the molecular correlates of cellular identity^9^. Over the past decade, thousands of intact, neurotypical human tissue samples from every major CNS region have been transcriptionally profiled with multiple technology platforms. These data provide an unprecedented opportunity to determine the core transcriptional features of cellular identity in the human CNS from the ‘top down’ by integrating cell-class-specific gene coexpression modules from a large number of independent datasets.

## RESULTS

### Gene coexpression analysis of synthetic brain samples accurately predicts differential expression among CNS cell classes

To illustrate the premise of our approach, we aggregated single-cell RNA-seq data from the adult human brain^1^ to create synthetic samples that mimic the heterogeneity of intact tissue (**Fig. 1A**). We performed unsupervised gene coexpression analysis of synthetic datasets and identified modules of coexpressed genes in each dataset that were maximally enriched with published markers^15, 16^ of astrocytes, oligodendrocytes, microglia, or neurons (‘cell-class modules’; e.g. **Fig. 1A**). Intuitively, the primary source of expression variation in a cell-class module is variation in the representation of that cell class in each sample. Mathematically, the vector that explains the greatest amount of expression variation in a coexpression module is its first principal component, or module 'eigengene' (**Fig. 1A**)^17^. This line of reasoning suggests that the eigengene of a cell-class module should approximate the relative abundance of that cell class in each sample. Because the precise cellular composition of each synthetic sample is known, we tested this hypothesis and found that actual cellular abundance was nearly indistinguishable from that predicted by cell-class module eigengenes (**Fig. S1A**).

**Fig. 1.**
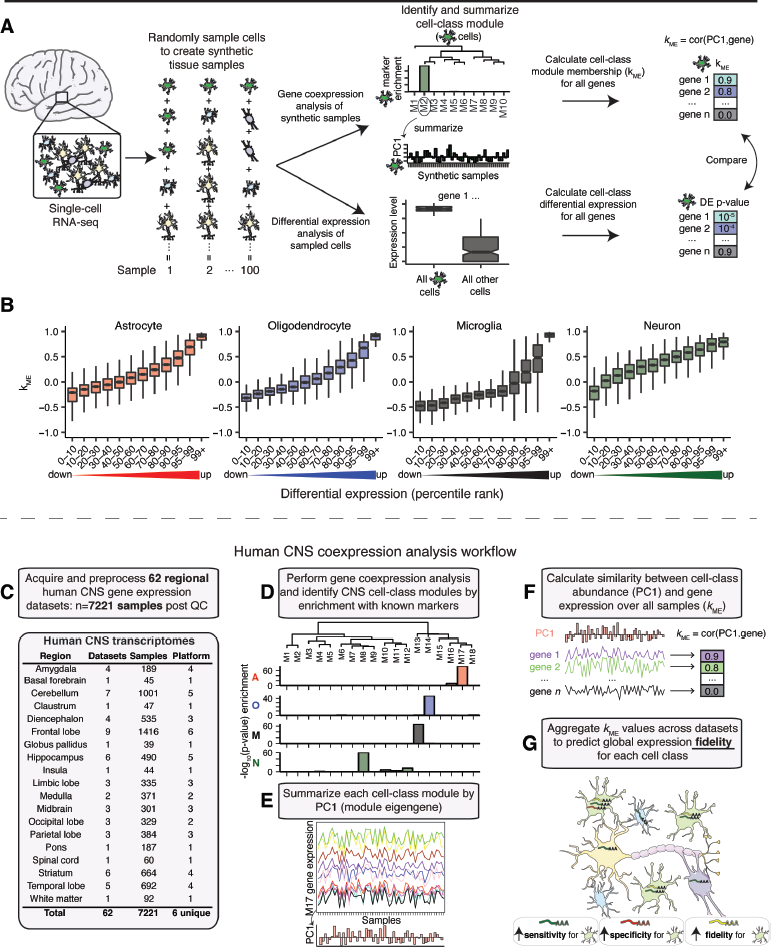
Rationale and workflow. **A)** Left: Single-cell RNA-seq data from adult human brain samples^1^ were randomly aggregated to create 100 synthetic tissue samples. Right (top): Unsupervised gene coexpression analysis of synthetic samples revealed CNS cell-class modules that were highly enriched with markers of astrocytes, oligodendrocytes, microglia, or neurons. Cell-class module membership strength (*k*_ME_) was calculated for all genes. Right (bottom): Using the same cells that were selected to create synthetic samples, single-cell differential expression analysis was performed for all genes with respect to each cell class. **B)** *k*_ME_ values for synthetic cell-class modules accurately predicted the results of differential expression analysis for each cell class (n=10 synthetic datasets; ‘up’ / ‘down’ denote up-and down-regulated genes for each cell class). **C)** 62 datasets representing diverse CNS regions and technology platforms were acquired and preprocessed (**Table S1**). **D)** Unsupervised gene coexpression analysis was performed for each dataset to identify modules of genes with similar expression patterns. Each module was summarized by PC1 (module eigengene). **E)** Published markers of each cell class were cross-referenced with all modules (Fisher’s exact test; **Table S2**). **F)** Cell-class module eigengenes were used to calculate the similarity between cellular abundance and genome-wide expression patterns (*k*_ME_) over all samples. **G)** Genome-wide *k*_ME_ values for significant cell-class modules were combined to yield a global measure of expression ‘fidelity’ for each gene with respect to each cell class. Schematic: A gene has high fidelity for a cell class if its expression is sensitive (it is consistently expressed by members of that cell class) and specific (it is not expressed by members of other cell classes).

To determine the affinity of each gene for each significant cell-class module, we calculated the Weighted Gene Coexpression Network Analysis measure of intramodular connectivity, or *k*_ME_^18^. *k*_ME_ is defined as the Pearson correlation between the expression pattern of a gene and a module eigengene. In the special situation of a cell-class module, *k*_ME_ therefore quantifies the similarity between the expression pattern of a gene and the relative abundance of that cell class in each sample. Because each sample is a heterogeneous mixture of cells, a high *k*_ME_ value for a cell-class module suggests that expression of the gene in that particular cell class is sensitive and specific. We tested this hypothesis by performing differential expression analysis of single-cell RNA-seq data for each cell class, restricting our analysis to exactly the same cells that were used to construct the synthetic brain samples. As shown in **Fig. 1B**, the genes that are most significantly up-regulated in a given cell class also have the highest *k*_ME_ values for the corresponding cell-class module. We obtained nearly identical results by aggregating single-cell RNA-seq data from the adult mouse brain^19^ (**Fig. S1B,C**). These findings demonstrate that gene coexpression analysis of intact CNS samples can determine which genes are most differentially expressed among CNS cell classes. More generally, our results suggest that it is not always necessary to physically isolate cells in order to ascertain their defining transcriptional features.

### Integrative gene coexpression analysis of intact tissue samples reveals consensus transcriptional profiles of major CNS cell classes in humans

To determine consensus transcriptional profiles of human CNS cell classes, we analyzed 7221 CNS transcriptomes from 840 neurotypical adult humans by combining data from eight studies^14,20-26^ and one resource (www.brainspan.org). These data were generated from intact postmortem tissue samples using diverse technology platforms (**Table S1**) and collectively represent billions of cells. Each sample was assigned to one of 19 broad neuroanatomical regions, resulting in 62 regional datasets (**Fig. 1C)**. After data preprocessing and quality control, each dataset consisted of ≥25 samples (median: 76) (**Table S1**). For each dataset, we performed unsupervised gene coexpression analysis and identified the module that was maximally enriched with published markers^15, 16^ of astrocytes, oligodendrocytes, microglia, or neurons (**Fig. 1D, Table S2**). PC1 of these modules was used to estimate the relative abundance of each cell class over all samples and calculate genome-wide *k*_ME_ values (**Fig. 1E,F**). Finally, we combined *k*_ME_ values for significant cell-class modules from all 62 datasets, producing a single value (z-score) for each gene that quantifies its global expression *fidelity* for each cell class (**Fig. 1G**). Importantly, estimates of fidelity were highly robust to the choice of gene set used for enrichment analysis (especially for glia; **Fig. S2**). Canonical markers consistently had high fidelity for the expected cell class and low fidelity for other cell classes (**Fig. 2A-D**). High-fidelity genes were also significantly and specifically enriched with expected cell-class markers from multiple independent studies (**Fig. 2A-D**). Compared to glia, the distribution of expression fidelity for neurons was compressed (**Fig. 2A-D**), likely reflecting neuronal heterogeneity among CNS regions. Genome-wide estimates of expression fidelity for major cell classes are provided in **Table S3** and on our web site (http://oldhamlab.ctec.ucsf.edu).

**Fig. 2.**
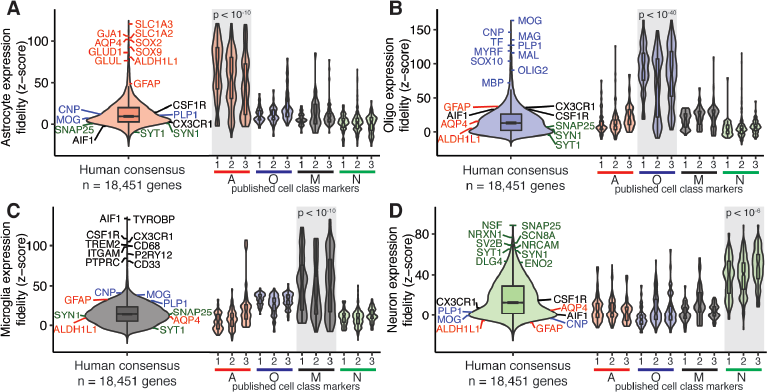
Integrative gene coexpression analysis of intact CNS transcriptomes reveals consensus transcriptional profiles of human astrocytes, oligodendrocytes, microglia, and neurons. **A-D)** Left: consensus gene expression fidelity distributions for human astrocytes (A), oligodendrocytes (O), microglia (M), and neurons (N). Canonical markers are labeled in red (A), blue (O), black (M), and green (N). Right: gene expression fidelity distributions for published sets of markers (A1-3, O1-3, M1-3, N1-3; Methods) were cross-referenced with high-fidelity genes (z-score >50). Gray shading: significant enrichment (Fisher's exact test).

### High-fidelity genes reveal the core transcriptional identities of major CNS cell classes in humans

The genes with the highest expression fidelity for major CNS cell classes are consistently coexpressed across regions and technology platforms (**Fig. S3**). This consistency suggests that high-fidelity genes can provide an unbiased view of the core transcriptional identities of major cell classes, thereby revealing novel cellular functions and biomarkers. We visualized the top 50 genes ranked by expression fidelity for each cell class to compare their expression levels, mutation intolerance, literature citations, cellular localization, and protein-protein interactions (PPI) (**Fig. 3A-D**). Overall, absolute expression levels of high-fidelity genes were highest for neurons and lowest for microglia (**Fig. 3A-D**, red tracks). However, for each cell class there was a wide range of expression levels for high-fidelity genes, suggesting parallel regulatory mechanisms and/or differential transcript stability.

**Fig. 3.**
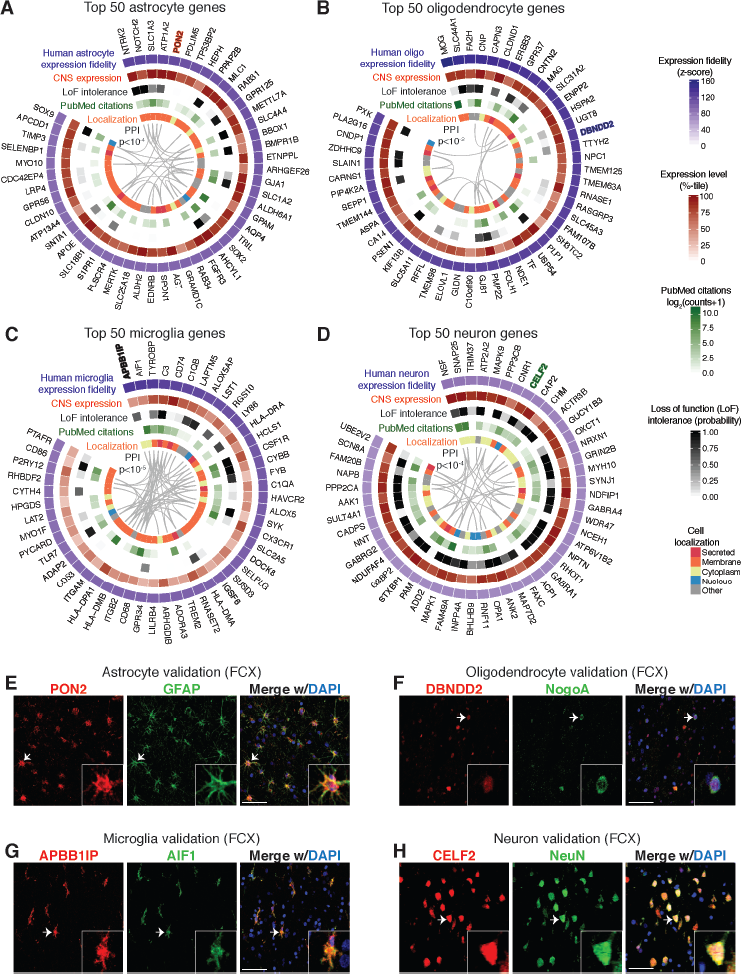
The core transcriptional identities of human astrocytes, oligodendrocytes, microglia, and neurons include known and novel biomarkers. **A-D)** The top 50 genes ranked by consensus expression fidelity for astrocytes, oligodendrocytes, microglia, or neurons. Expression levels: averages of mean percentile ranks for all datasets. Mutation intolerance: ExAC^27^. PubMed citations: queries for gene + cell class (e.g. gene symbol + 'astrocyte'). Cellular localization: COMPARTMENTS^28^. Predicted proteinprotein interactions (PPI): STRING^29^. Link=combined score >350. P-values for observed # links based on 100K random samples of 50 genes. **E-H)** Novel markers of human astrocytes (PON2), oligodendrocytes (DBNDD2), microglia (APBB1IP), and neurons (CELF2) in adult human dorsolateral prefrontal cortex (DLPFC; **E**: L5/6; **F**,**G**: white matter; **H**: L2/3). Arrowhead: cell in inset. Scale bar: 50pm.

To assess the tolerance of high-fidelity genes to loss-of-function (LoF) mutations, we analyzed data from the Exome Aggregation Consortium (ExAC), which summarizes the prevalence of coding mutations in ~61K human exomes^27^. Unexpectedly, high-fidelity neuronal genes were significantly less tolerant to LoF mutations than high-fidelity glial genes (**Fig. 3A-D**, black tracks). To determine whether high-fidelity genes have been studied in their respective cell classes, we searched PubMed for each gene symbol and the name of the cell class (**Fig. 3A-D**, green tracks). Interestingly, many searches returned no citations, highlighting critical gaps in our understanding of CNS cell biology. For example, the top microglial gene (amyloid beta precursor protein binding family B member 1 interacting protein, or *APBB1IP)* is unstudied in microglia.

We examined the cellular localization of proteins^28^ encoded by high-fidelity genes and observed another distinction between neurons and glia. Among the proteins encoded by genes in **Fig. 3A-D,** membrane localization was reported for 33 in astrocytes, 22 in oligodendrocytes, and 30 in microglia, but only 13 in neurons (inside track). This result may reflect the homeostatic functions of glia as sensors and regulators of extracellular CNS environments. More generally, the non-random distributions of cellular localizations suggest that high-fidelity genes are expressed at the protein level in the corresponding cell classes. To further explore this topic, we examined PPI^29^ among high-fidelity gene products for each cell class and observed significantly more interactions than expected by chance (**Fig. 3A-D**, interior lines).

Because high-fidelity genes should encode optimal biomarkers, we searched for high-fidelity genes in the Human Protein Atlas (http://www.proteinatlas.org) to identify novel reagents for labeling human CNS cell classes. We identified validated antibodies for PON2 (astrocytes), DBNDD2 (oligodendrocytes), APBB1IP (microglia), and CELF2 (neurons) (**Fig. 3A-D**). Dual immunostaining with canonical markers revealed almost perfect concordance in human frontal cortex (**Fig. 3E-H**).

### Gene coexpression analysis of intact tissue samples reveals the core transcriptional features of diverse CNS cell classes

Variation among intact tissue samples can also reveal transcriptional features of less abundant cell classes in the human CNS. Following the general strategy outlined in **Fig. 1**, we calculated genome-wide expression fidelity for human cholinergic neurons, midbrain dopaminergic neurons, endothelial cells, ependymal cells, choroid plexus cells, mural cells, oligodendrocyte progenitor cells, and Purkinje neurons (**Figs. 4, S4; Table S3**). This analysis correctly assigned high-fidelity scores for canonical markers of these cells. For example, choline acetyltransferase *(CHAT)*, the high-affinity choline transporter *(SLC5A7)*, and the vesicular acetylcholine transporter *(VACHT)* were all ranked within the top ~0.2% of all genes for cholinergic neuron expression fidelity, while claudin 5 *(CLDN5)*, tyrosine kinase with immunoglobulin like and EGF like domains 1 *(TIE1)*, and platelet and endothelial cell adhesion molecule 1 *(PECAM1)* were all ranked within the top ~0.3% of all genes for endothelial cell expression fidelity (**Table S3**). Comparisons with published gene sets revealed that high-fidelity genes were significantly and specifically enriched with expected markers of each cell class from multiple independent studies. Furthermore, novel markers predicted by our analysis were validated by *in situ* hybridization in the adult mouse brain^30^ (**Figs. 4, S4**).

**Fig. 4.**
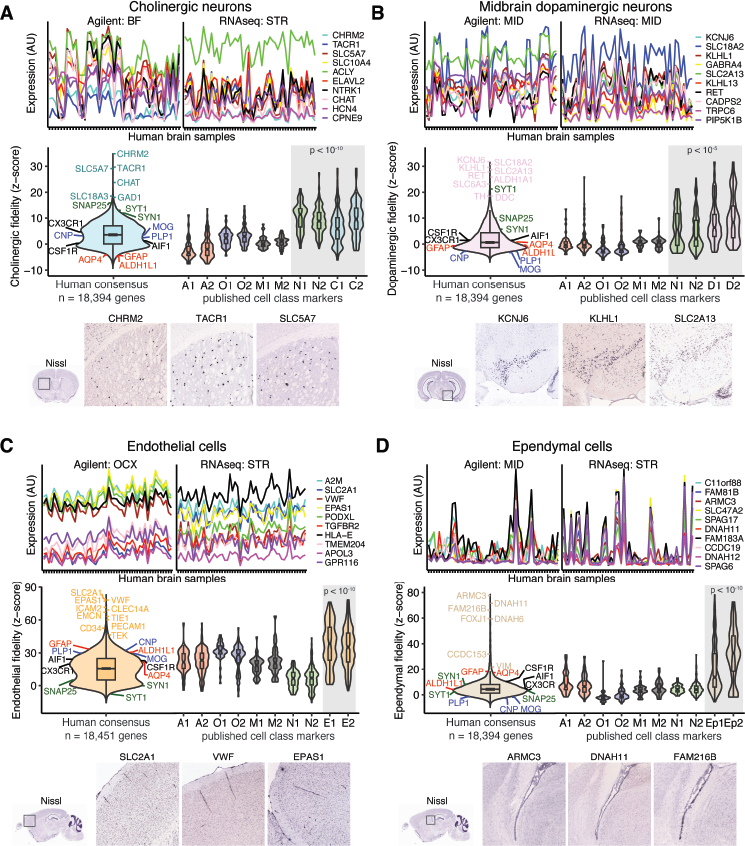
Variation among intact tissue samples reveals transcriptional signatures of human cholinergic neurons, midbrain dopaminergic neurons, endothelial cells, and ependymal cells. **A-D)** Top: high-fidelity genes for each cell class (top 10 are shown) are consistently coexpressed in independent datasets. Middle: consensus gene expression fidelity distributions for each cell class with canonical markers of major cell classes labeled in green (neurons), red (astrocytes), blue (oligodendrocytes), and black (microglia). Gene expression fidelity distributions for published sets of markers (Al, A2, O1, O2, M1, M2, N1, N2, C1, C2, D1, D2, E1, E2, Ep1, Ep2; Methods) were cross-referenced with high-fidelity genes (top 3 percentile). Gray shading: significant enrichment (Fisher's exact test). Bottom: mouse *in situ* hybridization data^30^ for high-fidelity genes in dorsal striatum (**A**), ventral midbrain (**B**), cortex (**C**), and lateral ventricle (**D**).

### High-fidelity genes enable predictive modeling of gene expression in transcriptomes from intact tissue samples

The reproducibility of gene coexpression modules corresponding to major cell classes (**Table S2, Fig. S3**) suggests that transcriptional variation among intact CNS samples can be modeled as a function of cellular abundance. We explored this topic systematically by performing multiple linear regression in 47 CNS datasets with ≥40 samples to determine how much expression variation in a shared set of ~9600 genes could be explained by variation in the abundance of neurons, astrocytes, oligodendrocytes, and microglia. To estimate the relative abundance of each cell class in each dataset, we summarized the expression patterns of high-fidelity genes (**Fig. 5A**). To avoid circularity, we used a leave-one-out cross-validation strategy to redefine high-fidelity genes for each dataset by recalculating expression fidelity for each cell class using the remaining 46 datasets (as in **Fig. 1C-G**). Prior to modeling, each dataset was downsampled (n=40) to facilitate comparisons of results; this process was performed iteratively to ensure robustness (**Fig. 5A**).

**Fig. 5.**
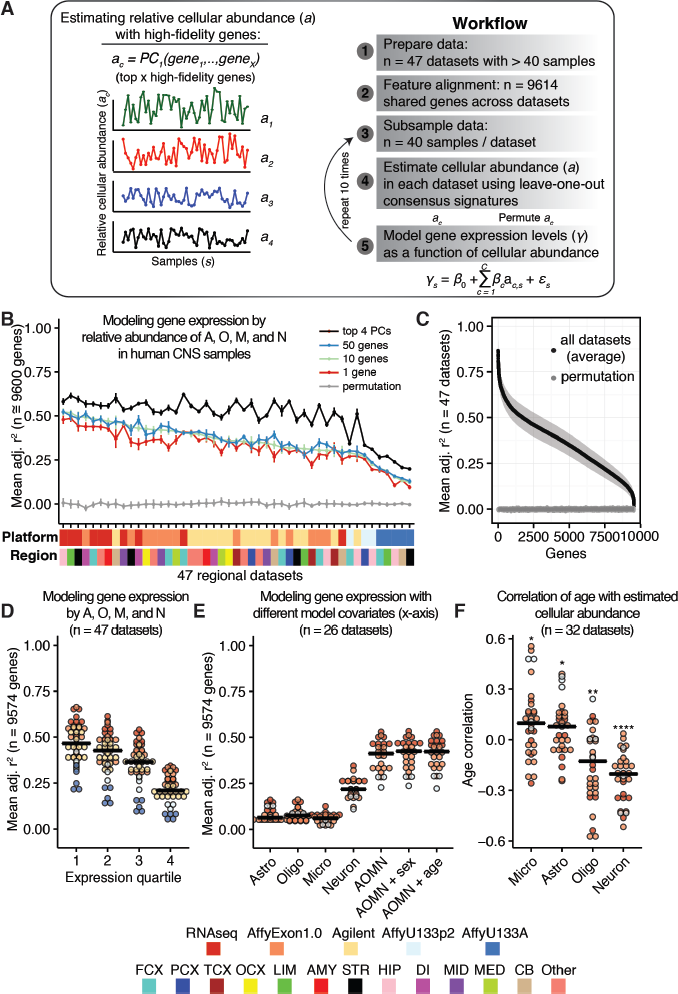
Variation in cellular abundance predicts gene expression in transcriptomes from intact CNS samples. **A)** Strategy for modeling gene expression in intact human CNS samples as a function of inferred cellular abundance. **B)** Total *%* variance explained (mean adj. r^2^) for ~9600 genes whose expression levels were modeled as a function of inferred astrocyte, oligodendrocyte, microglia, and neuron abundance in 47 datasets (subset to 40 samples; values are mean +/-2 s.e.m., 10 iterations). **C)** Mean adj. r^2^ values for individual genes from (**B**) over 47 datasets. Grey envelope: loess smoothed C.I. (+/-2 s.e.m., 10 iterations). **D)** Mean adj. r^2^ values for genes from (**B**) grouped by mean expression quartiles (each point is one dataset). **E)** Mean adj. r^2^ values for 7 different models (restricted to datasets w/ sex and age: GSE46706, GTEx, GSE11882, GSE25219). **F)** Pearson correlation of inferred cellular abundance with age (* p<0.05; ** p<0.01; **** p<0.0001, one-sample Wilcoxon signed-rank test). Horizontal bars (**D-F**): median; points colored by technology platform. Estimated cellular abundance

Implementing this strategy, we obtained several important results (**Fig. 5B**). First, using only one gene (with the highest fidelity) as a surrogate for each cell class, our models explained 32.2% of total transcriptional variation averaged over all datasets and up to ~50% in some datasets (vs. ~0.1% for permuted data). Second, increasing the number of gene surrogates/cell class (e.g. using the top 10 or top 50 high-fidelity genes) provided only modest performance improvements (unless otherwise stated, subsequent models used the top 10 high-fidelity genes). Third, prediction accuracy depended strongly on technology platform (p<10^-7^, ANOVA) but not CNS region (p=0.92, ANOVA). Among microarrays, older platforms fared substantially worse than newer platforms, while RNA-seq generally outperformed all microarrays.

Despite their simplicity, our models explained >50% of expression variation, averaged over all datasets, for ~2000 genes (**Fig. 5C**). Over all genes, the average amount of expression variation explained by our models followed a sigmoid function (**Fig. 5C**). We benchmarked model performance against the maximal explanatory power of any 4 predictors by using PC1-4 from each dataset as covariates for multiple regression. On average, PC1-4 explained 49.6% of total gene expression variation over all datasets (**Fig. 5B**). Thus, modeling gene expression in the human CNS as a function of neuron, astrocyte, oligodendrocyte, and microglia abundance achieved, on average, 72.0% of the maximal explanatory power for all datasets and 80.1% for RNA-seq datasets (**Fig. 5B**).

We reasoned that model performance for RNA-seq might exceed that for microarrays since the latter have many probes for transcripts that are unlikely to be expressed in the CNS. We therefore stratified genes by expression levels and examined model performance. As expected, predictive power decreased at lower expression levels, with the sharpest decline between the 3rd and 4th quartiles (**Fig. 5D**).

We next explored how transcriptional variation related to variation in the abundance of individual cell classes, sex, and age. We found that neuronal abundance explained more transcriptional variation than glial abundance (**Fig. 5E**). After controlling for variation in the abundance of major cell classes, model performance did not substantially improve by including sex or age as covariates (**Fig. 5E**). We further explored this topic by correlating the estimated abundance of each cell class with age in 32 CNS datasets. We found that neuronal and oligodendroglial abundance were negatively correlated with age, while astrocytic and microglial abundance were positively correlated (**Fig. 5F**). These results suggest that age-related changes in gene expression in bulk CNS transcriptomes are primarily driven by age-related changes in cellular composition.

### Gene expression modeling applications

The ability to predict gene expression in transcriptomes from intact CNS samples has substantial implications for many areas of neurobiological inquiry. We illustrate the relevance of this approach through comparative analysis of gene expression models in disease, among CNS regions, and between species.

### Application #1: Contextualizing disease genes and modeling gene expression in pathological samples

Using a curated database of results from genetic association studies^31^, we asked whether genes associated with CNS diseases are enriched among genes primarily expressed by astrocytes, oligodendrocytes, microglia, or neurons (**Fig. 6A-B)**. Clustering of select CNS diseases by enrichment p-values revealed several interesting findings. First, with the exception of ALS, genes associated with neurodegenerative disorders were most enriched among genes expressed by microglia and astrocytes. Second, genes associated with neurodevelopmental disorders, epilepsy, and psychiatric disorders were most enriched among genes expressed by astrocytes and neurons. Third, genes expressed by astrocytes consistently showed the greatest enrichment with candidate CNS disease genes.

Beyond broad associations between diseases and cell classes, gene expression modeling can also reveal which cell class is most likely to express a candidate disease gene. For example, we modeled gene expression for Alzheimer’s diseases (AD) risk genes as a function of neuronal, oligodendroglial, astrocytic, and microglial abundance in transcriptomes from intact neurotypical adult human temporal cortex (**Fig. 6C**). Expression levels of early-onset AD risk genes *APP* and *PSEN1* were mostly explained by variation in neuronal and oligodendroglial abundance, respectively. In contrast, expression levels of late-onset AD risk genes *APOE* and *TREM2* were mostly explained by variation in astrocytic and microglial abundance, respectively. These results were highly consistent across 47 CNS datasets (**Fig. 6D**).

**Fig. 6.**
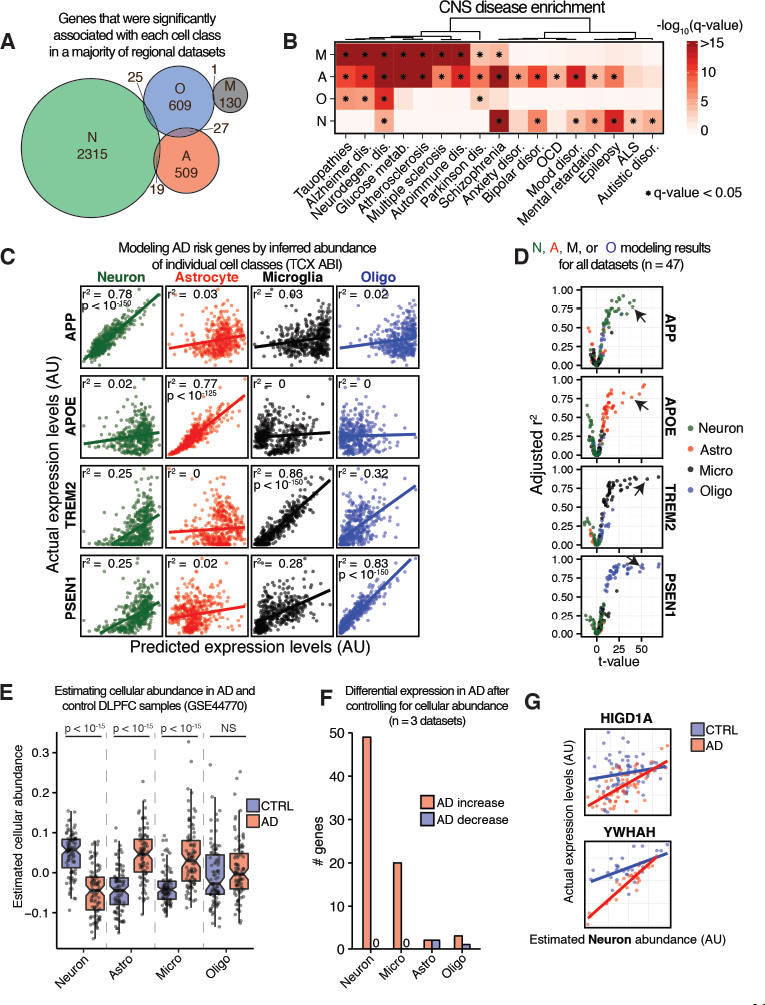
Gene expression modeling offers new avenues for studying human CNS diseases. **A)** *#* genes associated with each human cell class (p<8.37×10^−9^, Bonferroni correction for *#* gene models). **B)** Enrichment analysis (Fisher's exact test) of genes from (**A**) with human CNS disease genes from Phenopedia^31^. FDR-adjusted p-values (q-values) are shown^40^. **C)** Modeling results in human temporal cortex (TCX ABI; i.e. 1 dataset) for 4 AD risk genes. **D)** Modeling results for genes from (**C**) in 47 datasets (≥40 samples). **E)** Top 10 high-fidelity genes were used to estimate the relative abundance of neurons, astrocytes, microglia, and oligodendrocytes in DLPFC from control (CTRL) and AD^33^ (**Fig. 4A**). P-values: Wilcoxon rank-sum test. **F)** Gene expression modeling in 3 datasets^32-34^ reveals consistent cell-class-specific expression changes in AD after controlling for differences in cellular abundance (p<0.05 based on 1000 permutations of sample labels). **G**) Examples of two genes that are up-regulated in AD neurons (top^33^; bottom^34^).

Compared to control (CTRL) human brain samples, AD samples should contain fewer neurons and proportionately more glia. We tested this hypothesis by using expression patterns of high-fidelity genes to infer the relative abundance of neurons, astrocytes, microglia, and oligodendrocytes in 3 gene expression datasets from intact postmortem brain samples of CTRL and AD subjects^32-34^. We observed a highly significant decrease in neuronal abundance in AD in all datasets (**Figs. 6E**, **S5A-B**). In 2 out of 3 datasets, there were significant increases in the relative abundance of astrocytes and microglia in AD, with similar trends in the third (**Figs. 6E**, **S5A-B**). Interestingly, there were no significant differences in oligodendrocyte abundance between CTRL and AD in any dataset (**Figs. 6E**, **S5A-B**). This strategy can help determine whether variable cellular composition is associated with diverse CNS disorders.

Because AD brain samples tend to have fewer neurons and proportionately more astrocytes/microglia than CTRL, differential expression analysis of intact tissue samples will reveal down-regulation of neuronal transcripts and up-regulation of astrocytic/microglial transcripts. However, predictive modeling can identify cell-intrinsic transcriptional differences between CTRL and AD that are independent of changes in cellular composition. This strategy is analogous to that of Kuhn et al.^35^, except here we use expression patterns of high-fidelity genes to estimate cellular abundance. Surprisingly, after controlling for differences in cellular composition between CTRL and AD, we identified many genes that were consistently up-regulated in AD neurons (**Fig. 6F, Table S4**). These genes did not include canonical AD risk genes (**Fig. S5C**), but rather genes involved in protein ubiquitination, catabolism, proteasome degradation, and mitochondrial function (**Fig. S5D**), suggesting efforts by AD neurons to mitigate the effects of misfolded protein aggregates. Examples are shown in Figs. 6G, **S5**.

### Application #2: Identifying transcriptional differences in major cell classes among CNS regions

We recalculated expression fidelity separately for each CNS region with ≥3 datasets and performed hierarchical clustering for each cell class (**Fig. 7A-D**). Regional differences in expression fidelity were greatest for neurons, with a clear bifurcation between cortical/subcortical structures (**Fig. 7D-E**). In contrast, expression fidelity for oligodendrocytes was very similar among brain regions (**Fig. 7B,E**). Comparatively, microglia and astrocytes exhibited more regional variation in expression fidelity than oligodendrocytes, but less than neurons (**Fig. 7A,C,E**).

**Fig. 7.**
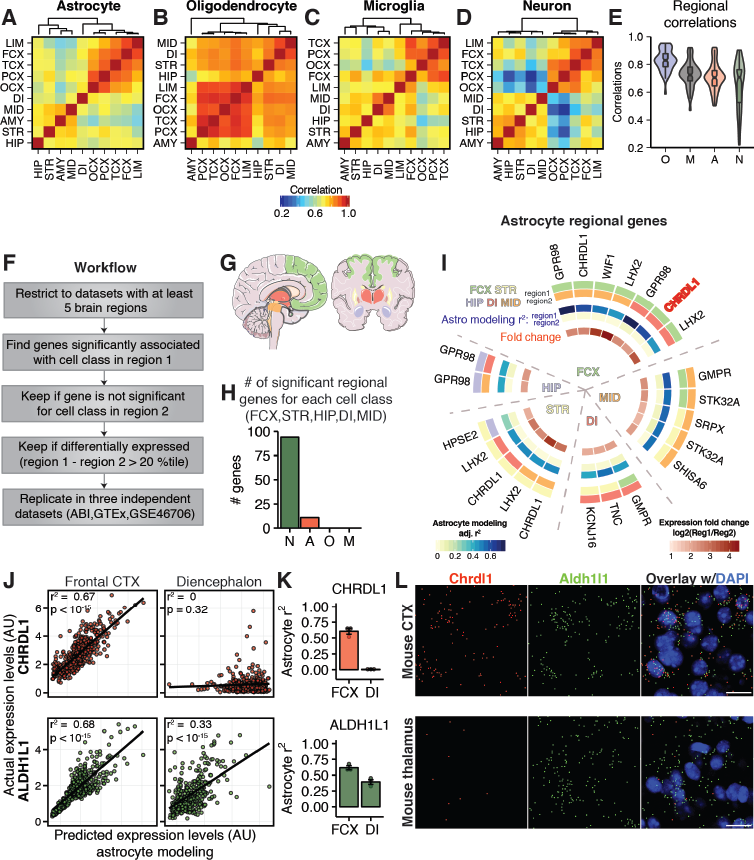
Regional expression fidelity and predictive modeling reveal astrocyte heterogeneity in the human brain. **(A-D)** Hierarchical clustering of human brain regions (excluding cerebellum) based on Pearson correlations among regional expression fidelity for each cell class (n=18451 genes, ≥3 datasets/region). **E)** Distributions of correlations in (**A-D**). **F)** Workflow to predict regional expression dif-ferences in specific cell classes. Significance threshold: p<2.67×10^−8^ (Bonferroni correction for total # of gene models). **G)** Analyzed brain regions: frontal cortex (FCX), striatum (STR), hippocampus (HIP), diencephalon (DI), and midbrain (MID). **H)** Total # of region-specific genes conservatively predicted for each cell class. **I)** Genes predicted to be expressed by human astrocytes in restricted brain regions. **J)** Modeling of *CHRDL1* and *ALDH1L1* (+ control) as a function of inferred astrocyte abundance in example datasets (FCX/DI from ABI). **K)** Modeling results for same genes in 3 datasets (ABI, GTEx, and GSE46706). **L)** Single-molecule FISH of *Chrdll* and *Aldhlll* in mature mouse brain (P30). Scale bar: 20µm.

We developed a conservative strategy to identify binary expression differences in major cell classes among human brain regions (**Fig. 7F-G, Table S5**). Using these criteria, many genes were predicted to distinguish regional subpopulations of neurons (**Figs. 7H, S6**). Using the same criteria, we found no evidence for binary expression differences among regional subpopulations of microglia or oligodendrocytes (**Fig. 7H**). However, we did predict binary expression differences among regional subpopulations of human astrocytes (**Fig. 7H-I**). For example, *CHRDL1* was predicted to be expressed by astrocytes in frontal cortex and striatum, but not by astrocytes in diencephalon and midbrain (**Fig. 7I-K**). To validate this prediction, we performed single-molecule fluorescent *in situ* hybridization (FISH) for *Chrdl1* and *Aldh1l1* in cortical and thalamic samples from mice. *Aldh1l1* is expressed ubiquitously by astrocytes^15^ and was detected in mouse cortex and thalamus (**Fig. 7J-L**). Expression of *Chrdl1* colocalized with *Aldh1l1* in mouse cortex but not thalamus (**Fig. 7L**), as predicted.

### Application #3: Identifying transcriptional differences in major CNS cell classes between species

We analyzed 1346 mouse brain transcriptomes to determine genome-wide expression fidelity for astrocytes, oligodendrocytes, microglia, and neurons (**Tables S1**, **S6**; **Fig. S7**). Over all homologous genes, expression fidelity was significantly correlated between mice and humans for each cell class, with the greatest similarity for neurons (**Fig. 8A**). We note that the strong conservation of neuronal expression fidelity relative to glia is mirrored at the protein level: high-fidelity neuronal genes are significantly less tolerant to LoF mutations than high-fidelity glial genes (**Fig. 3A-D**, black tracks). These findings may indicate that neurons are under greater evolutionary constraint than glia.

**Fig. 8.**
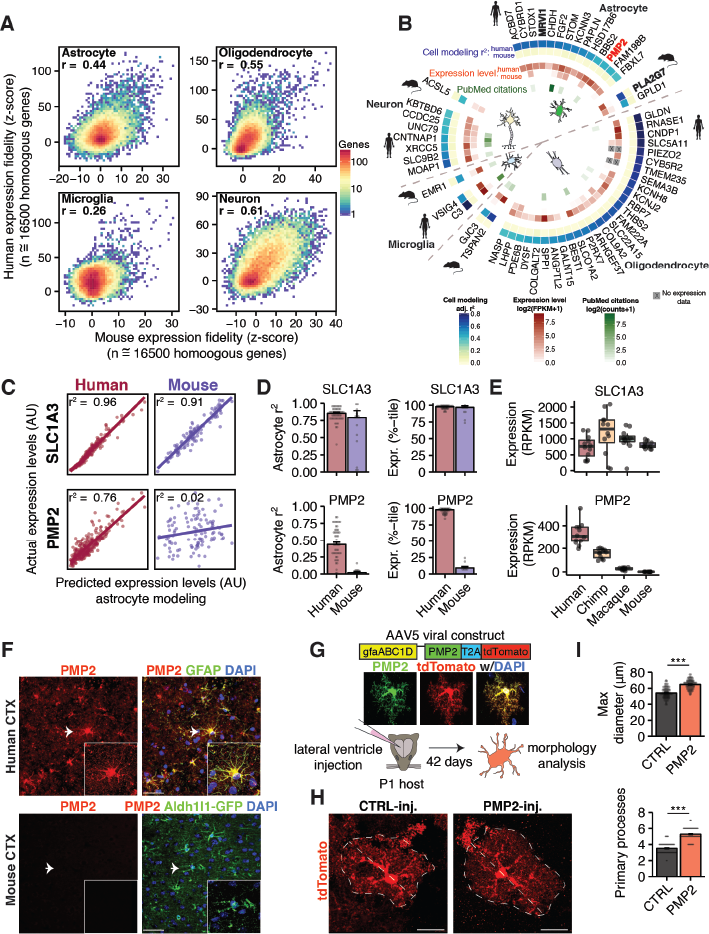
Gene expression modeling identifies cell-class-specific transcriptional differences between humans and mice. **A)** Comparison of gene expression fidelity in humans and mice for each cell class. **B)** Predicted cell-class-specific transcriptional differences between humans and mice. Expression levels are from independent datasets^3, 41^ that were not used to predict species differences. PubMed citations obtained as in **Fig. 3**. **C)** Example modeling results in humans (Hs.PCX.ABI) and mice (Ms.GSE64398) (**Table S1**). *SLC1A3* is expressed by astrocytes in both species and *PMP2* by astrocytes in humans but not mice. **D)** Astrocyte modeling results and mean expression percentiles for genes in (**C**) from all datasets. Error bars: s.e.m. **E)** *SLC1A3* and *PMP2* expression in human, chimpanzee, macaque, and mouse prefrontal cortex^38^. **F)** Immunostaining for PMP2 in adult human DLPFC and P42 mouse neocortex. Arrowheads: cells in insets. Scale bar: 40µm. **G)** Experimental strategy for studying *PMP2* effects on mouse astrocytes. **H)** Representative examples of CTRL and PMP2-infected astrocytes in mouse neocortex. Dashed lines outline cell and max diameter through nucleus. Scale bar: 20µm. **I)** Quantification of max diameter and # of primary processes in CTRL and PMP2-infected astrocytes. n=4 animals/group, n>15 astrocytes/animal, mean ± s.e.m., Welch’s t-test, *** p<0.001.

We applied stringent criteria and identified 50 genes predicted to be ‘on’ in human CNS cell classes and ‘off in the corresponding mouse CNS cell classes (**Fig. 8B, Table S7**). Only 6 genes were predicted with the opposite pattern (**Fig. 8B, Table S7**), which may reflect the smaller number of mouse transcriptomes analyzed. ~85% of predicted transcriptional differences between humans and mice were in glia (**Fig. 8B**). Because these differences could reflect an evolutionary gain of expression in one species or loss in the other, we analyzed 476 outgroup samples from chimpanzee and macaque brains (**Table S1**). Of the 50 genes predicted to be expressed in human but not mouse cell classes, 29 were significantly associated with the same cell class in at least one primate dataset; conversely, of the 6 genes with the opposite pattern, none was significantly associated with the same cell class in any primate dataset (**Table S7**). For example, expression variation of *MRVI1* was largely explained by variation in astrocyte abundance in primates, but not mice (**Figs. 8B, S8A-B**). Conversely, expression variation of *PLA2G7* was largely explained by variation in astrocyte abundance in mice, but not primates (**Figs. 8B, S8A-B**). Single-molecule FISH in human and mouse cerebral cortex confirmed that expression of *MRVI1* and *PLA2G7* is specific to human and mouse astrocytes, respectively (**Fig. S8C-D**).

To provide proof of concept for the ability of our analyses to deliver functional insights into the unique biology of human brains, we focused on a major unexplained cellular phenotype, which is the fact that human astrocytes are much larger than mouse astrocytes (as well as non-human primate astrocytes)^36^. This phenotype has important implications for neuronal function, since the domain of one human astrocyte can encompass up to ~2MM synapses vs. only ~100K synapses for one mouse astrocyte^36^. We reasoned that genes expressed by human but not mouse astrocytes might contribute to this phenotype. We were particularly intrigued by peripheral myelin protein 2 (PMP2; **Fig. 8B**), which encodes a fatty-acid binding protein made by Schwann cells that is important for maintaining membrane lipid composition^37^. In the human CNS, expression of *PMP2* was extremely high (mean percentile: 96.2) and largely explained by variation in astrocyte abundance, while in the mouse CNS expression of *PMP2* was effectively absent (mean percentile: 11.2) and unrelated to variation in astrocyte abundance (**Fig. 8B-D**). Furthermore, independent RNA-seq data from human, chimpanzee, macaque, and mouse prefrontal cortex^38^ revealed a monotonic increase in *PMP2* expression from mouse to human (**Fig. 8E**).

Immunostaining showed widespread PMP2 in human neocortical astrocytes (**Fig. 8F**). In contrast, PMP2 was undetectable in mouse neocortex (**Fig. 8F**), despite robust expression by Schwann cells (**Fig. S8E**). To test whether PMP2 could increase mouse astrocyte size *in vivo*, we delivered a viral construct expressing *PMP2* under an astrocyte-specific promoter to neonatal mouse brains and analyzed the morphology of transduced astrocytes after 42d (**Fig. 8G**). Forced expression of *PMP2* in mouse astrocytes significantly increased their maximum diameter and number of primary processes (**Fig. 8H-I**). The increase in máximum diameter corresponded to an increase in mouse astrocyte volume of ~50% (assuming sphericity). To further validate this finding, we repeated the experiment with a different viral construct and obtained nearly identical results (**Fig. S8F**). To our knowledge, these data provide the first molecular explanation for morphological differences between human and mouse astrocytes. More generally, our findings illustrate how variation among intact tissue samples can predict cell-class-specific transcriptional features with important functional implications for human neurobiology.

## DISCUSSION

We have described a novel, ‘top-down’ approach to reveal the core transcriptional features of cellular identity via integrative gene coexpression analysis of intact tissue samples. Compared to ‘bottom-up’ methods such as FACS, IP, and SC/SN, the main advantages of our approach are as follows: i) elimination of the need for fresh tissue; ii) applicability to huge amounts of existing data; iii) elimination of technical variability caused by tissue dissociation and cDNA preamplification; iv) elimination of sampling bias associated with cell/nucleus capture; and v) ability to derive highly robust inferences about the core transcriptional features of cellular identity based on aggregate analysis of billions of cells.

Our approach also has important limitations. False-positive associations can result from technical factors such as batch effects or biological factors such as cellular collinearity. For example, we consistently observed that genes with high expression fidelity for oligodendrocytes had higher expression fidelity for microglia (and vice versa) than they did for astrocytes or neurons. Because oligodendrocytes and microglia are more abundant in white matter than gray matter^39^, variation in the ratio of white matter to gray matter in CNS samples drives covariation in the abundance of these cell classes and the genes that they express. False-negative associations can result from technical factors such as limitations in dynamic range/transcriptome coverage or probe failures, as well as biological factors such as alternative splicing. Notwithstanding these limitations, the genes with the highest expression fidelity for major CNS cell classes are already remarkably stable.

It is interesting to consider the ability of our approach to detect transcriptional signatures of less abundant cell classes (e.g. **Figs. 4, S4**). The ability to discern a gene coexpression signature of a cell class in transcriptomes from intact tissue samples depends on many factors, including its representation, the uniqueness and abundance of its transcripts, its stoichiometry with other cell types, the technology platform, the algorithmic approach, and the sampling strategy^8^. Some of these factors can be optimized to improve sensitivity. Ultimately, however, we envision future studies that combine the benefits of top-down and bottom-up strategies to fully deconstruct the transcriptional architecture of biological systems.

Our estimates of gene expression fidelity for major cell classes were highly robust to the choice of gene set used for enrichment analysis, but more so for glia than neurons. This result indicates that neuronal diversity may require additional strategies to optimize estimates of neuronal expression fidelity, particularly on a regional basis. For example, the neuronal gene sets used in this study do not capture the transcriptional profile of cerebellar granule neurons, which is highly distinct^14,22,26^. To better account for neuronal diversity, future studies may utilize additional neuron subtype-specific or composite gene sets for enrichment analyses.

Our results suggest that the functional identity of a cell class can be conceived as a vector of genes ranked by the fidelity with which they are expressed in that cell class relative to all other cells in the biological system of interest. An advantage of this framing is that it is inherently context-dependent. Beyond revealing novel biomarkers and cellular phenotypes, such definitions can provide ‘molecular rulers’ for measuring the validity of human cells derived *in vitro* for disease modeling and cell replacement therapies. In addition, these definitions can be tested in *de novo* CNS transcriptomes for their ability to predict gene expression levels through mathematical modeling.

Multivariate analyses of CNS transcriptomes often use module detection/clustering methods or projection methods such as principal component analysis. Although these methods have produced many important insights, they are inherently descriptive and do not lend themselves easily to comparisons among independent datasets. Because the building block of any biological system is the cell, and cells are distinguished by the genes that they express, an alternative approach is to model expression levels of individual genes as a function of variation in cellular composition. We have shown how expression patterns of high-fidelity genes can be used as covariates in multiple linear regression models for this purpose. The resulting models are grounded in biology, easily compared among independent datasets, and capable of extracting cell-class-specific insights from intact tissue samples. Using this approach, we explored how predictive models of gene expression in transcriptomes from intact CNS samples can inform studies of aging, disease genes, pathological samples, regional heterogeneity, and species differences. We elaborate upon our findings in the Supplementary Discussion.

The analyses presented in this study are based on a simple idea: variation in cellular composition among intact tissue samples will drive covariation of transcripts that are uniquely or predominantly expressed in specific kinds of cells. Although we have focused here on gene expression, our approach can also be applied to other types of molecular data, thereby offering a generalizable strategy for determining the core molecular features of cellular identity in intact biological systems.

## ACKNOWLEDGMENTS

We are grateful to Brad Dispensa, Joe Hesse, Dirk Kleinhesselink, and Jason Jed for technical support. We thank Annette Molinaro for statistical consultations, David Rowitch for astrocyte discussions, and Eric Huang and Mercedes Paredes for human brain samples. Due to space limitations, we apologize that many relevant and important publications are not cited. This work was supported by the UCSF Program for Breakthrough Biomedical Research (MCO), which is funded in part by the Sandler Foundation, a Scholar Award from the UCSF Weill Institute for Neurosciences (MCO), and NIMH R01MH113896 (MCO).

## AUTHOR CONTRIBUTIONS

KWK and MCO conceived and designed the analytical strategies and wrote the manuscript. KWK performed most data analyses and histological experiments. KWK and HO performed *PMP2* expression experiments under supervision from AVM.

